# Emergence of evolutionary stable communities through eco-evolutionary tunneling

**DOI:** 10.1101/271015

**Authors:** Seyfullah Enes Kotil, Kalin Vetsigian

**Author notes:** **Address correspondence to** Kalin Vetsigian.

## Abstract

Ecological and evolutionary dynamics of communities are inexorably intertwined. The ecological state determines the fate of newly arising mutants, and mutations that increase in frequency can reshape the ecological dynamics. Evolutionary game theory and its extensions within adaptive dynamics (AD) have been the mathematical frameworks for understanding this interplay, leading to notions such as Evolutionary Stable States (ESS) in which no mutations are favored, and evolutionary branching points near which the population diversifies. A central assumption behind these theoretical treatments has been that mutations are rare so that the ecological dynamics has time to equilibrate after every mutation. A fundamental question is whether qualitatively new phenomena can arise when mutations are frequent. Here we describe an adaptive diversification process that robustly leads to complex ESS, despite the fact that such communities are unreachable through a step-by-step evolutionary process. Rather, the system as a whole tunnels between collective states over a short time scale. The tunneling rate is a sharply increasing function of the rate with which mutations arise in the population. This makes the emergence of ESS communities virtually impossible in small populations, but generic in large ones. Moreover, communities emerging through this process can spatially spread as single replication units that outcompete other communities. Overall, this work provides a qualitatively new mechanism for adaptive diversification and shows that complex structures can generically evolve even when no step-by-step evolutionary path exists.

## Main Text

Understanding the consequences of the feedback between ecological and evolutionary dynamics is a fundamental challenge^1–6^, particularly in asexual microbial communities, where new strains with altered ecological interactions can readily arise through mutational events such as replication errors and horizontal gene transfer^7–14^. In the absence of mutations, the strain abundance dynamics can be represented as a non-linear dynamical system with fixed dimensionality determined by the number of strains^15–18^, which allows us to understand and classify long-term community composition in terms of dynamical attractors. It is far less clear, however, how to analyze the dynamics when mutations stochastically introduce new dimensions and ecological interactions^19–22^.

Important progress was made with the advent of Evolutionary Game Theory^23–26^, which established the notion of evolutionary stable states as long-term endpoints of the eco-evolutionary dynamics. An evolutionary stable state is a community that has reached an ecological attractor that cannot be invaded by any possible mutant. Therefore, an evolutionary stable community (ESC) would persist indefinitely despite mutations. These ideas were further elaborated on through the framework of AD^27–29^, which went beyond analysis of equilibria. Here community evolution is represented as a sequence of mutational events (Fig 1a). After every mutational event the ecological dynamics has time to equilibrate and reach a new ecological attractor. Thus, the evolutionary dynamics consists of a series of stochastic jumps from one ecological attractor to another, and an ecological attractor can in general transition to many different attractors through different mutations (Fig 1b). The space of possible strains and attractors is typically continuous, reflecting the continuous nature of phenotypes. It was recognized that some ESS can be unreachable by evolution (“Gardens of Eden”)^30^, and therefore the notion of evolutionary convergence was introduced to indicate ecological attractors towards which all nearby attractors evolve. Strikingly, evolutionary stability and evolutionary convergence are independent notions that can occur in all possible combinations^27,31^. States that are both convergent and stable are end-points of the eco-evolutionary dynamics. In contrast, ecological attractors that are evolutionary convergent but unstable (branching points) lead to adaptive diversification events, where a lineage splits into multiple coexisting lineages. Thus, AD has provided tools for analyzing equilibrium end-points of the eco-evolutionary dynamics and adaptive diversifications^32^.

A fundamental limitation of the AD framework is the assumption that ecologically relevant mutations are rare, so that the ecological dynamics has time to equilibrate between mutations^28^. In a sense, it is assumed that there is a separation of time scales between the ecological and evolutionary dynamics. However, even if the rate of discovery of ecologically relevant mutations is low per individual, many communities contain very large numbers of individuals. Therefore, it is quite possible that a new mutant will start invading before the transient dynamics triggered by the previous successful mutant has expired. This would be especially true if the selection coefficients of invading mutants are low, so that the transient ecological dynamics is long.

An important question, then, is whether the realistic regime of a finite mutation rate per population, rather than an infinitesimal one, can exhibit qualitatively new behaviors that fall outside the scope of adaptive dynamics. Recently, we computationally examined the eco-evolutionary dynamics in microbial communities in which bacteria consume a single resource in a two-dimensional environment and compete through interactions mediated by antibiotic production and degradation^33^. Each strain is characterized by its costly investment in antibiotic production or degradation with respect to each antibiotic, and mutants with different levels of antibiotic production and degradation can arise (Fig. 1cde, Methods). We found that, for a broad range of parameters, ESCs with 3-strains and 5-strains robustly arise in simulations with one and two antibiotics, respectively, when starting with a single strain. Thus, stable communities maintained solely by antibiotic-mediated interactions can spontaneously evolve. Here, we investigate the mechanism behind these adaptive diversifications. We show that, while locally convergent, the ESCs are globally inaccessible through step-by-step adaptive dynamics, starting from simple single-strain populations (Fig. 1b). The robust emergence of these “forbidden” ESCs is only possible when the timescales of the ecological and evolutionary dynamics interfere with each other through a process we term “eco-evolutionary tunneling”.

**Fig 1.**
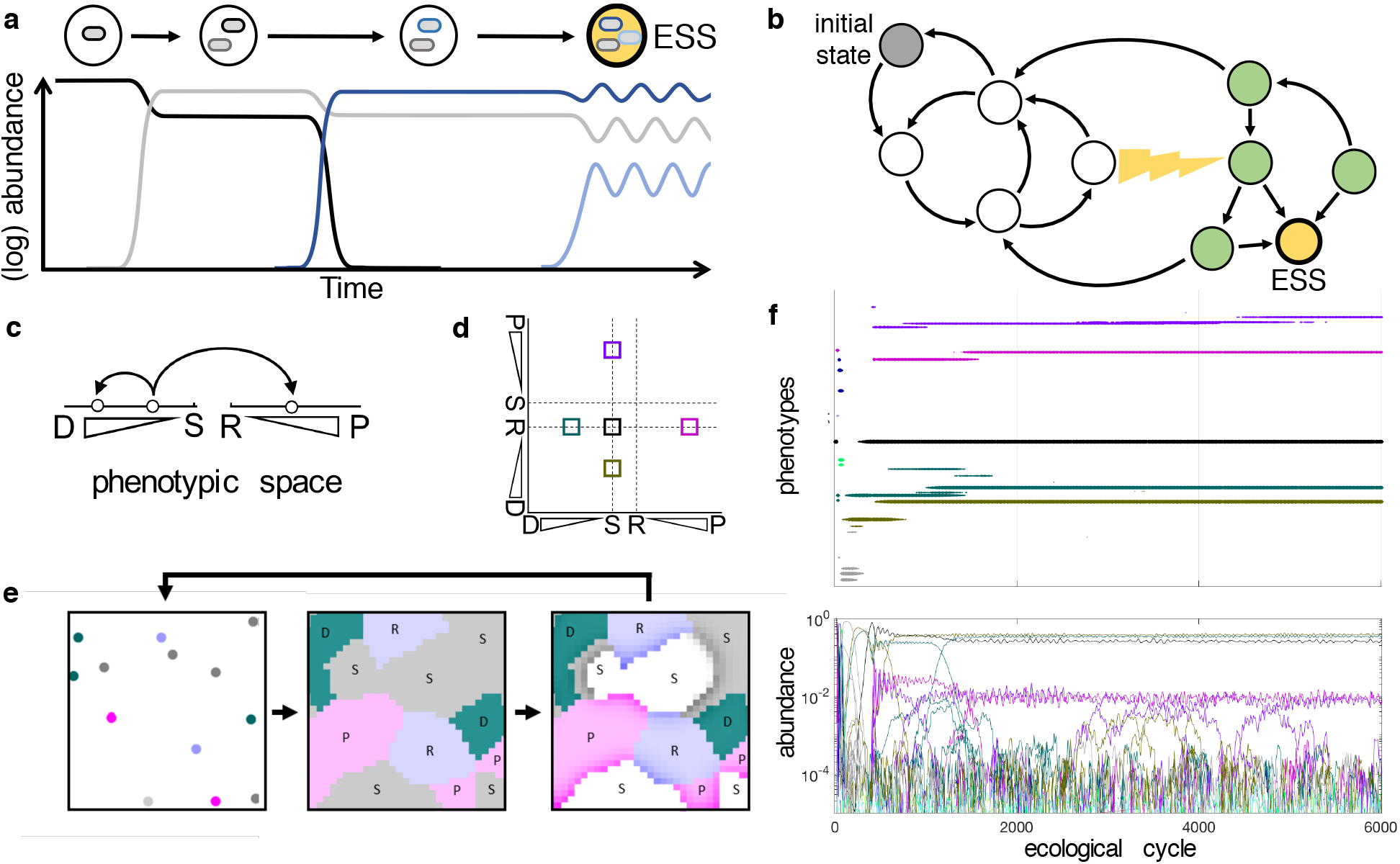
Evolution of antibiotic interactions leads to robust emergence of ESCs that are unreachable through adaptive dynamics. **(a)** Schematic of AD. Mutant invasions (arrows) lead to transitions between ecologically stable communities (circles) containing one or more strains. ESS corresponds to a state from which no further transitions are possible (yellow circle). The abundances of different strains over time (colored lines) corresponding to the transitions are also shown. **(b)** A more general AD graph in which multiple paths are possible, and in which certain ecologically stable communities (green) are unreachable starting from an initial community (grey). These communities can, however, be reached through eco-evolutionary tunneling that falls outside the scope of AD (yellow lightning). **(c)** In simulations, strains genetically encode different levels of antibiotic production (P) or degradation (D). They can also be sensitive to antibiotics (S) or be resistant via an antibiotic efflux (R). New strains can emerge through mutations (arrows). **(d)** The space of possible strain phenotypes in simulations with two antibiotics. The 5-strain ESC that emerges in (f) is indicated by colored squares. **(e)** Simulations of microbial communities proceed though ecological cycles. During each cycle, spores of different strains (different colors) are scattered on a 2D plane and grow as colonies until all space is filled. Colonies then produce and degrade diffusing antibiotics according to their phenotypes. Some regions die (white), while sporulation is enhanced if neighbors are inhibited (darker colors). **(f)** In a simulation with two antibiotics a 5-strain ESC robustly arises. (Top) Shown are existing phenotypes (projected into 1D) as a function of time. (Bottom) Shown is the abundance of different strains over time. Different colors correspond to different phenotypic classes (Methods).

ESCs appear to come together all at once (Fig. 1f). To disentangle the sequence of events leading to their formation, we lowered the mutation rate per population in the hope of stretching out the assembly process over time. We found that while this tends to dramatically delay the emergence, the stable community still snaps together on a very short timescale (Fig. 2a). Before the transition, the system spends a lot of time in a regime of rapid turnover (RT), in which the dominant strains are continuously outcompeted by new mutants. This regime ends abruptly with the emergence of a persistent multi-strain community that can then evolve towards the convergent ESC. This transition is stochastic, and the wait time (*T*) is exponentially distributed (Fig. 2b, Supplementary Fig. 1, Supplementary Fig. 2). This suggests a simple two-state representation in which the RT state is metastable and decays into the ESC state by stochastically tunneling through a barrier with a constant probability per unit time (Fig. 2c).

We next investigated how the tunneling rate (*r*) depends on the mutation rate (*μ*) and population size (*N*). If the community emerges step-by-step (Fig 1a), we expect that, at low *μN*, the inverse of the mean community formation time would be simply proportional to the mutation rate per population (*μN*), independent of ecological details (Methods). In sharp contrast, we found that *r* is proportional to (*μN*)^2^ for the 3-strain community that emerges in a universe with one antibiotic, and *r* exhibits a complex dependence with an approximate (*μN*)^3.7^ scaling for the 5-strain community that emerges in a universe with two antibiotics (Fig. 2d, Supplementary Fig. 3). The superlinear dependence of *r* clearly demonstrates that ESC emergence cannot be understood within the context of adaptive dynamics and, strikingly, that emergence becomes ever more efficient at high mutation rates or population sizes – requiring fewer mutational events. Furthermore, the different scaling exponents at low *μN* in the two examined cases indicate that there are different classes and orders of eco-evolutionary emergence.

**Fig 2.**
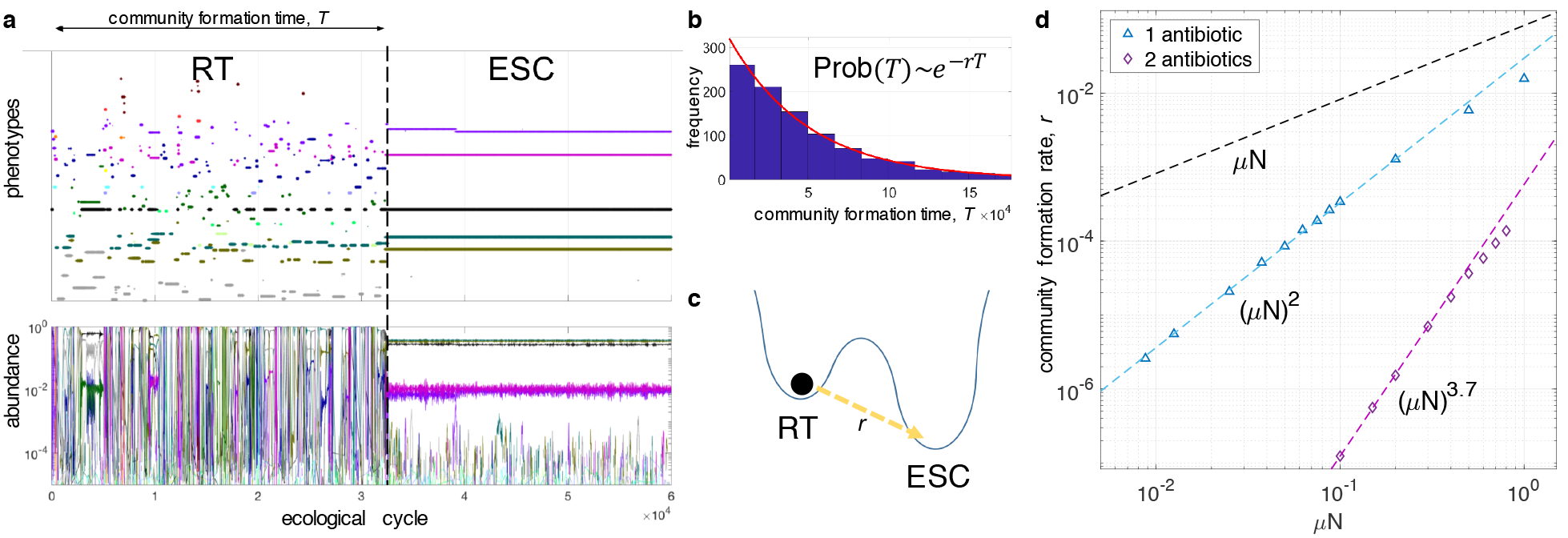
ESCs emerge abruptly through eco-evolutionary tunneling whose rate scales superlinearly with mutation rate and population size. **(a)** At lower mutation rates, the dynamics can spend a considerable time in a rapid strain turnover phase (RT) before suddenly arriving at a stable solution (ESC). **(b)** The community formation time is exponentially distributed, indicating a constant probability of formation per unit time. **(c)** A two-state representation of the system in which a metastable RT state decays into an ESC state by tunneling through a barrier with a constant probability per unit time, *r*. **(d)** The community assembly rate *r*, as computed from thousands of replicate simulations (Supplementary Fig. 1, Supplementary Fig. 2), is shown as a function of *μ* at fixed *N* for simulations with one antibiotic (blue triangles) and two antibiotics (purple diamonds). Note the logarithmic scale on both axes. Straight lines (dashed, blue and purple) are fitted to the data to demonstrate power law dependence at low *μN*. The expected slope from AD for *r*=1/<*T*> is shown as a reference (dashed black line). The estimation errors of *r* are smaller than the symbols. By independently varying *μ* and *N*, we verified that the controlling parameter is the product *μN*, which is the mutation rate per population (Supplementary Fig. 3).

Community emergence can be understood as a probabilistic escape from the RT regime. This can happen when the turnover dynamics passes through an exit window and requires a well-timed sequence of mutant invasions, which we term a high-order eco-evolutionary event (Fig. 3). The emergence of a 3-strain community maintained by one antibiotic provides the simplest possible example and requires a single well-timed mutation that happens during an ongoing transition of the RT dynamics from one state to another (Fig. 3a). Given an ongoing transition, the probability (*p*) of such a mutation arising is proportional to (*μN*)*W* for small *p*, where *W* is the length of the window of opportunity. On the other hand, the frequency (*f*) with which a system passes through an exit window is proportional to *μN*. Therefore, *r* = *fp* ~(*μN*)^2^, explaining our empirical finding for simulations with one antibiotic.

Simulations with two antibiotics provide a more general example in which multiple well-timed mutant invasions are required, and in which a community can emerge through multiple possible pathways (Fig. 3bc, Supplementary Fig. 4, Supplementary Fig. 5). The pathway on Fig. 3b shows that two higher-order eco-evolutionary events, separated by a stable stepping-stone community, can act in combination to enable complex community formation. A simple analysis suggests that each higher-order event increases the scaling exponent of *r* based on the number of well-timed mutations, while the number of AD steps does not affect the scaling (Methods). This analysis predicts *r* ~(*μN*)^4^, which is close but not identical to the observed exponent of about 3.7. The discrepancy might be due to the fact that we have not reached the true asymptotic behavior at low *μN*. Alternatively, the non-trivial scaling exponent might be an emergent property of the collective dynamics.

Strikingly, the most frequently utilized pathway goes directly from one strain to five strains, without any ecologically stable intermediates (Fig. 3c). This would appear to imply *r* ~(*μN*)^5^, making the dominance of this pathway puzzling (Methods). The resolution of this paradox lies within the continuous nature of the phenotypes and the resulting possibility that ecologically distinct strains can happen to be close in fitness in the absence of other strains. Such combinations of strains can persist for a long time and effectively act as if they were ecologically stable stepping stones, thus reducing the expected emergence order (Methods). Thus, near neutral coexistence of strains can provide a path for efficient ESC formation.

**Fig 3.**
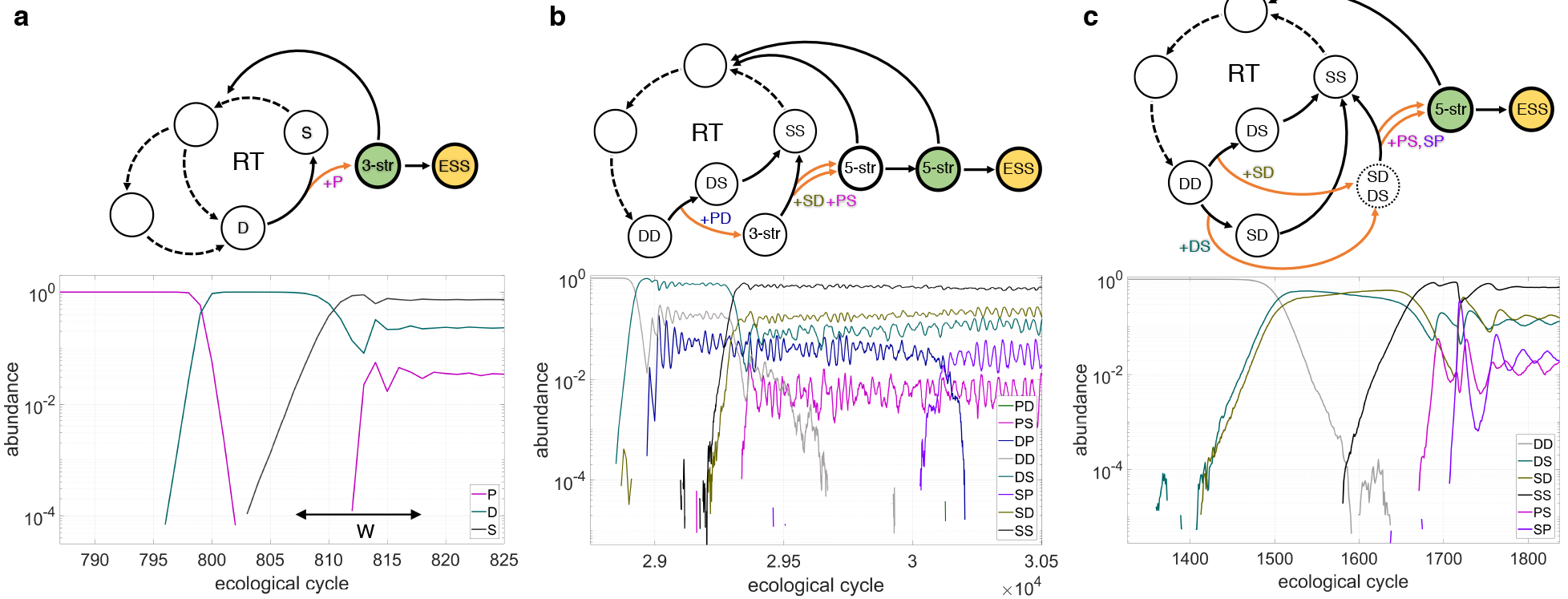
Pathways of ESC emergence. Shown are sample simulation trajectories leading to multi-strain community formation (bottom) together with the corresponding state transition diagrams (top). Higher-order eco-evolutionary events are indicated as orange arrows. Communities of the same type as ESC (ESC-like, Methods) are shown in green, **(a)** In simulations with one antibiotic, the strain turnover dynamics occasionally leads to a situation in which a resident degrader strain (D) is being outcompeted by a sensitive strain (S). As S increases in frequency, antibiotic producer strains (P) become evolutionary favored as well. However, if a producer strain arises too late, it will simply outcompete the sensitive strain. A producer must arise within a particular time-window (W) for an ecologically stable 3-strain community to form, if such a community is possible at all given the particular numerical values of the P and D phenotypes, **(b)** Shown is one of the possible pathways of formation of a 5-strain community maintained by two antibiotics. The turnover dynamics leads to a state dominated by a strain that degrades both antibiotics (DD). While this strain is being outcompeted by a strain that degrades only the first antibiotic (DS), a PD-strain arises and invades as well, leading to a formation of a 3-strain (DP, DD, and DS) ecologically stable community (see Methods for two-letter strain nomenclature). During the invasion of this community by a strain sensitive to both antibiotics (SS), two additional mutants (SD and PS) also invade, which leads to an ecologically stable 5-strain community (after the loss of the DD-strain). **(c)** Shown is the most frequently utilized pathway for 5-strain community formation. A DD-strain is outcompeted by DS and SD strains. Through clonal interference, this two-strain community (dotted line circle) persists long enough for the SS-strain to arise. As SS increases in frequency, antibiotic producer strains become selected for (SP and PS). If these strains arise before the extinction of DS or SD, an ecologically stable 5-strain community can form.

We next investigated the emergence of ESC in spatially extended systems. Instead of having one community that is well-mixed at the end of every ecological cycle, we simulated communities connected by migration. We examined the rate of community formation as a function of the migration rate (Fig. 4a). The limit of low migration corresponds to many independent small communities, whereas very high migration corresponds to a single, large community that exhibits a higher rate of community emergence due to the non-trivial *μN* scaling. Strikingly though, community formation is fastest at intermediate migration rates – indicating that spatial structure facilitates eco-evolutionary emergence. A way to understand this is that, at low enough migration rates, the communities at different locations fail to synchronize their rapid turnover dynamics, and therefore immigration can introduce new strains into a community, effectively augmenting mutation.

Another spatial phenomenon we might expect from systems with a stable and a meta-stable phase is that the stable phase will grow when in contact with the metastable one (Fig. 4b). This is known as nucleated growth^34^. Therefore, we asked whether a small spatial region containing an ESC would expand and outcompete neighboring communities. This is indeed what happens (Fig. 4c). This is significant because it constitutes an example of a multi-strain community that grows and spreads as a self-replicating unit. Thus, we have shown that eco-evolutionary tunneling can lead to the spontaneous formation of higher-level, self-replicating composites of strains. The ability to spread implies that even if the emergence of a diverse community is unlikely at any given location, it only needs to arise once in order to become ecologically dominant.

The formation of stable communities through a stochastic escape from a regime of rapid strain turnover via several well-timed mutant invasions is a mechanism for adaptive diversification that is fundamentally different than that relying on branching points. Its hallmarks - abrupt emergence without a complete set of stable intermediates, exponential distribution of the community formation time, superlinear scaling of formation rate with population size, colonization ability - are all experimentally observable, either directly from time-series data of individual trajectories (as illustrated by Fig. 3) or through replicate microcosm experiments.

This mechanism demonstrates that evolution of complex communities that is impossible through independent evolutionary steps can nevertheless be efficient and generic in sufficiently large communities. It constitutes an example of an emergent phenomenon that is fundamentally eco-evolutionary - it is only possible when the timescales of the ecological and evolutionary dynamics overlap. The mutation rate per population is a parameter that controls this overlap and can be viewed as a measure of evolutionary “temperature,” with AD being akin to a zero-temperature limit. In analogy with communities of atoms and molecules where numerous phenomena and phases of matter (e.g. liquids) only exist at non-zero temperature, we expect that there is a rich spectrum of eco-evolutionary phenomena and states that are only possible in the regime in which population mutation rates are not close to zero and multiple ecologically relevant invasions can be underway at the same time. We anticipate that further work will elucidate many additional such phenomena in both sexual and asexual populations.

**Fig 4.**
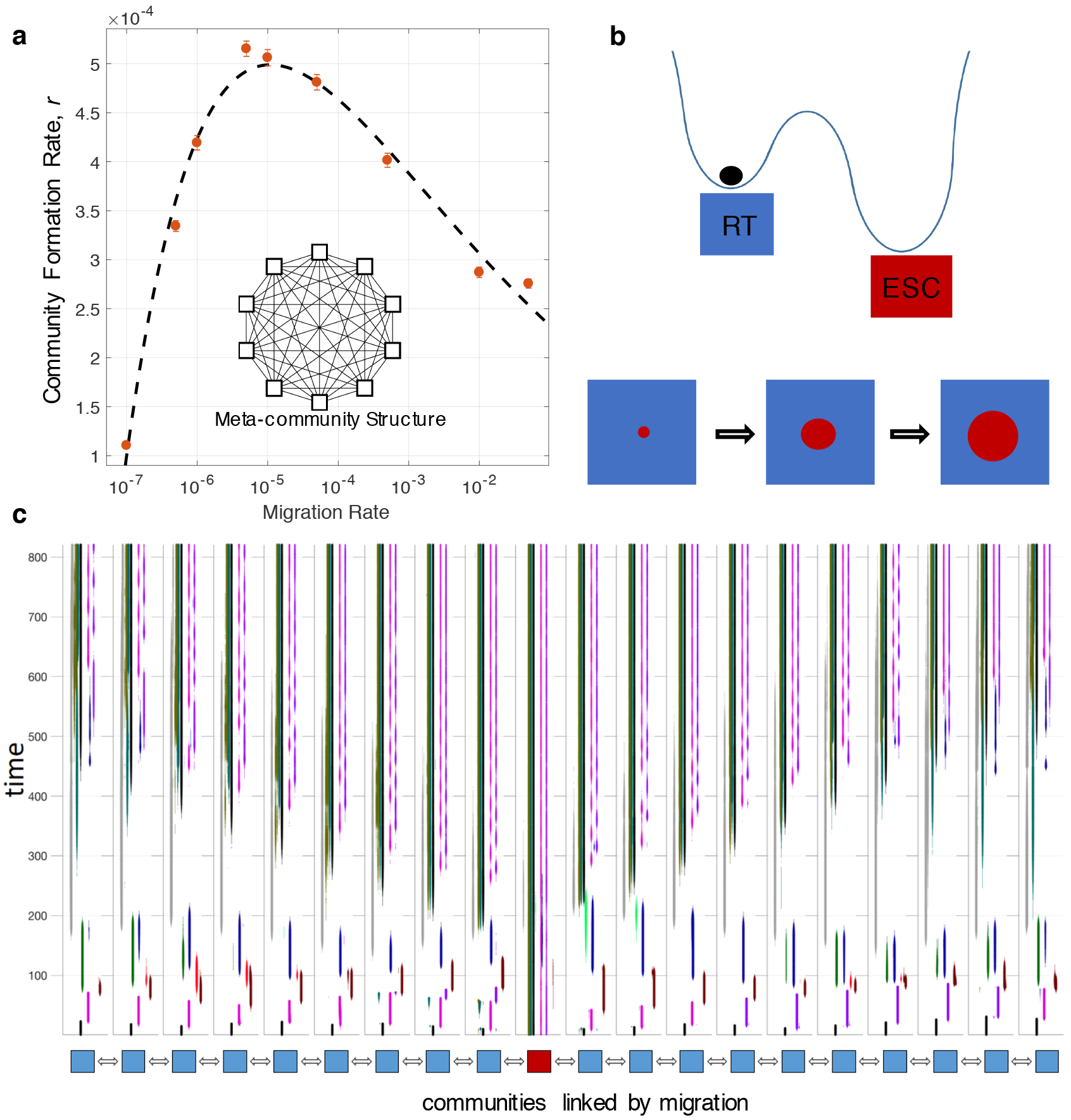
Accelerated emergence and spread of ESC in spatially extended systems. **(a)** The community formation rate *r* is shown as a function of the migration rate for 10 patches linked by migration. The rate is highest at intermediate migration rates, where the spatial structure is non-trivial. **(b)** For systems with a stable (maroon) and a meta-stable (blue) phase, nucleated growth is expected in which a (sufficiently large) seed of the stable phase grows and displaces the meta-stable phase. **(c)** A one-dimensional chain of patches linked by migration is simulated in a universe with two antibiotics. The central patch is seeded with the 5-strain ESC and the other patches contain a single strain sensitive to both antibiotics. Shown for each patch are the existing phenotypes as a function of time. The 5-strain pattern corresponding to ESC spreads to neighboring communities at an approximately constant speed, after an initial delay due to the special initial state of the non-central patches.

The emergence of complex Nash equilibria in which multiple strategies stably coexist is of interest not only in ecology but also in economics. While economic theory often assumes that systems are already at Nash equilibria^35^, recent work has highlighted the difficulties of reaching them^36^. The ideas developed here suggest that the near-simultaneous spread of multiple new beneficial strategies among players might facilitate the establishment of Nash equilibria.

## ACKNOWLEDGEMENTS

This work was supported by the Simons Foundation, Targeted Grant in the Mathematical Modeling of Living Systems Award 342039, the National Science Foundation Grant DEB 1457518. This research was performed using the compute resources and assistance of the UW-Madison Center for High Throughput Computing (CHTC) in the Department of Computer Sciences. We thank Simon A. Levin for insightful discussions.

## AUTHOR CONTRIBUTIONS

SEK and KV designed the study, analyzed the data, performed the simulations, and wrote the manuscript.

## Methods

The computer simulations used in this study are based on the model described in^33^ and are briefly summarized on Fig.1cde. For the results presented on Fig. 4, the model was extended to include multiple patches linked by migration as described below. All the simulations and numerical analyses were performed in MATLAB using custom scripts that are available upon request.

*μ* is the probability of mutation per individual per ecological cycle. *N* is the number of spores/individuals at the beginning of each ecological cycle and is kept constant by randomly sampling *N* spores at the end of the ecological cycle to start a new one. Mutations happen between ecological cycles, and the number of mutations that happen in the population between cycles is Poisson distributed with mean *μN*. The relative probabilities of different types of mutations are kept fixed when varying *μ* or *N* and are as described before.

### Definition of ecological and evolutionary stability

*Ecological stability* is defined as the long-term persistence of diversity when mutations are turned off (despite demographic noise coming from finite population sizes). *Evolutionary stable communities* (ESC) are ecologically stable communities for which no mutants exist that can increase in frequency and alter the community composition.

### Classification of strains into discrete phenotypic classes

While phenotypes are continuous in simulations, we found it useful for analysis, visualization, and interpretation of results to assign them to several discrete categories. With respect to a single antibiotic we distinguish D, S, R, and P strains. A D-strain is one that has the ability to degrade the antibiotic at some level. A P-strain is a strain that produces the antibiotic at a level that can inhibit a neighboring S-strain colony. A R-strain has an efflux pump making it resistant to the antibiotic (just as a P-strain) but does not produce enough antibiotics to inhibit neighbors. The S-strain does not degrade, produce or efflux antibiotics and is therefore sensitive to the antibiotic. In simulations with two antibiotics we have 16 classes that correspond to all possible pairs of {D, S, R, P}. For example, a DP-strain is a strain that can degrade the first antibiotic at some level and produce the second antibiotic at some level. On figures presenting evolutionary trajectories, strains from different phenotypic classes are shown in different colors, and strains from the same phenotypic class have the same color.

### ESC-like communities

In simulations, the true ESC is never reached as the selection coefficients of mutations that bring the community closer to ESC become vanishingly small, and the approach slows down. Furthermore, the gradual final approach to ESC proceeds through adaptive dynamics (selective sweeps within emergent ecotypes) and is not of primary interest. What we focus on is the transition from a population dominated by a single strain to an ecologically stable multi-strain community from which the evolution towards ESC can proceed entirely through adaptive dynamics. Correspondingly, we define the notion of an *ESC-like* community as is an ecologically stable community that consists of strains that belong to the same phenotypic classes (see above) as the ESC.

In simulations with 1 antibiotic, ESC consisted of 3-strains for the parameters we used: the S strain, a particular P strain, and a particular D strain. An ESC-like community, therefore, is an ecologically stable community consisting of a P-strain, a D-strain and the S-strain. In simulations with two antibiotics, ESC consists of 5-strains: particular PS, SP, DS, SD strains and the SS strain. An ESC-like community is an ecologically stable community containing 5-strains that belong to these 5 phenotypic classes.

### Determining the community formation time *T*

For each simulation, we determined the ecological cycle number (time) of the first occurrence (if any) of an ESC-like community. This is done algorithmically through the following heuristic procedure:

First, we determine the abundance over time for the ESC-like phenotypic classes over time. In simulations with 1 antibiotic, we keep track of the abundance of P, D and S phenotypic classes over time. For simulations with 2 antibiotics, we record the abundance of PS, SP, DS, SD and SS classes over time. For example, the abundance of the SP class is determined by summing the abundances of all SP-strains that exist at a particular time. Typically, the abundance was recorded every 10 ecological cycles.

To determine if an ESC-like community emerges at time-point *t*, we analyze the phenotypic class abundance trajectories over the 5 subsequent time-points (including the current time-point), spanning 50 ecological cycles. If the abundance of any ESC-like phenotypic class reaches zero during this interval, time-point *t* does not mark the onset of an ESC-like community. If all abundances stay positive, we compute the linear least-squares fits for each phenotypic class. We then calculate the relative slope of each linear fit by dividing the change in abundance by the maximum value over the time interval. If all the relative slopes (for the ESC-like phenotypes) are below 0.05, we mark *t* as the time of ESC-like community formation. For each simulation trajectory, the time of first emergence of an ESC-like community is taken to be the community formation time *T*.

The accuracy of this algorithm has been validated by manually examining thousands of evolutionary trajectories. The scaling of *r* with *μN* is insensitive to the time-interval and threshold used, though they might lead to a shift of the distribution of *T* by a constant amount or a multiplicative shift of *r* (e.g. due to rejection of very short-lived ESC-like communities).

### Verifying that the community formation time is exponentially distributed and determining the community formation rate *r*

For particular values of *μ* and *N*, thousands of replicate trajectories lasting for tens of thousands of ecological cycles (see Parameters for exact values) were generated (starting with different random number generator seeds). For each trajectory, we determined the community formation time *T*. If no community formed, we set T to infinity.

We then constructed the empirical cumulative distribution function CDF(*t*) = Prob(*T*<*t*), which specifies the fraction of trajectories with *T* smaller than *t*. If *T* is exponentially distributed we expect Prob(*T* ≥ *t*) = 1−CDF(*t*) = exp(−*rt*), and correspondingly a linear relationship between *y*=log(1−CDF(*t*)) and *t*. Strong linearity was indeed observed as shown on Supplementary Figures 1 and 2.

The slope provides an estimate of *r*. Note that the exponential distribution will be in general shifted due to the non-random initial condition and the finite duration of a higher-order eco-evolutionary event (See *τ*_0_ below). Correspondingly, we allowed the fitted line not to pass through the origin (*y* = −*rt* + *a*).

### Migration

A meta-community model of *H* patches is characterized by an *H*×*H* migration matrix *M*. *M*_i,j_ specifies the fraction of the population from community *j* that migrates to community *i* in one ecological cycle.

We considered two spatial settings in this paper:

1. All-to-all connected patches (Fig. 4a), for which we set *M*_i,j_ = *m*(1−δ_i,j_) + (1−(*H*−1)*m*)δ_i,j_, where δ_i,j_ is the Kronecker delta symbol. *H*=10 was used.
2. A ring of patches (Fig. 4c), implemented as *M*_i,j_ = (1−2*m*)δ_i,j_ + *m*(δ_i,j-1_+δ_i,j+1_+δ_i,H_ δ_j,1_+δ_i,1_ δ_j,H_). The last two terms connect patches 1 and *H* to create a ring. *H*=20 was used.

If *Y*_*jS*_ is the abundance of strain *s* in patch *j* at the end of an ecological cycle, we set 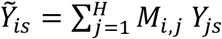, and start the next ecological cycle of patch *i* by randomly assigning strain phenotypes to *N* individuals according to a multinomial distribution with 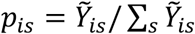. The ecological cycles of all patches are synchronous.

For the all-to-all connected meta-community, the overall community formation time was defined as the time when any of the communities reaches an ESC-like state according to the algorithm described above.

### The rate (1/<T>) of reaching a state through adaptive dynamics is *μN*

Adaptive dynamics assumes that the system is mutation limited. The system spends almost all of its time waiting for the next successful mutant to invade and drive its transition from one ecological attractor to another. The waiting times at each state (circle on Fig. 1ab) are exponentially distributed, and each possible transition (arrow on Fig. 1ab) is characterized by a transition rate - the probability per unit time that the adaptive mutant underlying the specific transition will appear and escape stochastic loss. Each transition rate is, therefore, proportional to *μN*. If the RT dynamics is driven by mutant invasions with non-negligible selection coefficients, then at low *μN* the transitions themselves will have negligible duration compared with the mean wait times. Correspondingly, the mean time it takes to hop along any particular sequence of state transitions is equal to the sum of mean waiting times along the trajectory. Therefore, the mean time to reach any state (ecological attractor) from any other is proportional to (*μN*)^−1^ (as long as no nearly neutral transitions are required). Based on the observation that the turnover of strains is indeed rapid, it is clear that the turnover dynamics is pushed forward by mutants with non-negligible selection coefficients.

### Rate of community emergence in simulations with one antibiotic

The basic idea is that the rapid turnover dynamics passes through exit windows. During such passes, there is some probability that a community will form though a higher-order eco-evolutionary event.

Let *τ* be the average time it takes the rapid turnover adaptive dynamics to return to an exit window, if currently at one. Based on the argument above, *τ* ~ (*μN*)^−1^ at low *μN*.

Let *p* be the probability that a community will form through a higher-order eco-evolutionary event during an ongoing passage through an exit window. In general, the dynamics will pass through an exit-window many times before eventually a community successfully forms. Let *n* = 1, 2, 3,… be the number of times the dynamics passed through an exit window before success. *n* is a geometrically distributed random variable, and <n> = 1/*p*. Correspondingly, the mean community formation time is:

<T> = (<n> − 1) *τ* + τ_0_ = *τ* (1−*p*)/*p* + *τ*_0_, where τ_0_ is the sum of the mean time to reach the exit window from the initial condition and the mean time it takes for the community to come together after reaching the exit window for the last time.

As *p* → 0, <*T*> → τ / *p*, and *r* → *p* / *τ*.

In the one antibiotic case, escape from the RT regime requires a single well-timed mutant invasion - a P-strain must invade during an adaptive dynamic transition from an S-strain dominated community to a D-strain dominated community. The P-strain must arise during some time window W; if the P-strain arises too late, it will take over the community dominated by the S-strain. Let *q* ~ *μN* be the probability per ecological cycle that such a mutant will arise and escape stochastic loss. Then *p* = 1 − (1 − *q*)^*W*^ ≈ 1 − exp(−*qW*). Therefore, *p* ~ *μN* for small *μN*, and correspondingly, *r* ~ (*μN*)^2^.

Notice that if *qW* is not small, the approximation *p* ~ *μN* no longer holds. In particular, as we increase *μN*, we would expect deviations from *r* ~ (*μN*)^2^. If *qW* is of order unity, then *p* can be close to 1 as well, and we might get *r* ~ *μN*. Thus, the scaling exponent of *r* can decrease at high enough μN. Deviations from power law scaling are indeed noticeable at high *μN* on Fig. 2d.

### Rate of community emergence in simulations with two antibiotics

Generalizing the logic from above, if *k* mutations need to arise during an ongoing adaptive dynamic transition for a community to emerge, this will lead to *p* ~ (*μN*)^*k*^ for small *μN*.

The pathway presented on Fig. 3b can be split into two tunneling transitions. First, a higher-order eco-evolutionary event is required to arrive at the 3-strain (SD+DP+DD) ecologically stable community. Repeating the argument for one antibiotic, the rate of arriving at this 3-strain community is *r*_3_ ~ (*μN*)^2^. The 3-strain community can be considered a part of an extended strain turnover loop. The formation of a 5-strain community requires a second higher-order eco-evolutionary event to exit the extended loop. During an invasion of the 3-strain community by a DS strain, two additional mutants (SS, SP) need to arise. Correspondingly the probability of success is *p*_3→5_ ~ (*μN*)^2^. Repeating the logic from the one antibiotic case, now with the extended turnover loop that incorporates the 3-strain community, we get: *r* = *r*_3_*p*_3→5_, and therefor *r* ~ (*μN*)^4^ at low *μN*.

For the pathway presented on Fig. 3c, 4 mutations need to arise during an ongoing transition from DD to, say, DS, which implies *p* ~ (*μN*)^4^ and, using *r* = *p*/τ, *r* ~ (*μN*)^5^. Thus, it would appear that this channel would be less likely than the other one. This indeed might be the case at very low *μN*. However, starting from the lowest *μN* levels we could computationally explore, this was one of the main channels for formation of 5-strain communities (Supplementary Fig. 5). A qualitatively new feature of this pathway is the DS+SD two-strain intermediate. The relative selection coefficients of the DS and SD strains depend on the properties of the DD strain they derive and take over from. Each time the turnover dynamics reaches a DD, it will have a somewhat different phenotype. Crucially, some DD strains will give rise to DS and SD strains that have nearly identical selection coefficients. If that happens DS and SD will coexist in a near neutral fashion for extended periods of time (clonal interference), virtually guaranteeing that this two-strain combination will persist until a mutant invasion. Thus, DS+SD can effectively act as a regular ecologically stable community awaiting a mutant invasion. The 5-strain community formation can, then, be viewed as a set of two higher-order eco-evolutionary events: the first creating the DS+SD state by means of an SD invasion during a DD to DS transition, and the second - creating the 5-strain community through PS and SP invasions during a DS+SD to SS transition. The rate of reaching SD+DS would be (μN)^2^, as for the one antibiotic case, and the probability of the second eco-evolutionary tunneling is (*μN*)^2^ because of the two required well-timed mutations. Therefore, we expect *r* ~ (*μN*)^4^ for pathways with long-lived SD+DS states, which is of the same order as the pathway on Fig. 3b.

### Parameters

The parameters are defined in^33^. The 2D Gaussian filter used in all steps was size 5×5 and σ = 1. α = 0.5, λ = 0.5, Γ = 0.3, δ = 2, *T*_p_ = 40, *T*_r_ = 10, *N* = 40, *L* = 100, *S* = 3600, *D*_max_ = 1000, *P*_max_ = 1000, μ = 10, {π_LF_, π_l_, π_g_} = {1/3, 1/3, 1/3}, ε = 0.

**Figure 1d:** C_D_^0^ = 0.03, C_D_ = 4×10^−4^, C_P_ = 4×10^−4^, C_R_ = 0.3, μN = 4, N = 1.44×10^5^.

**Figure 2a:** C_D_^0^ = 0.03, C_D_ = 4×10^−4^, C_P_ = 4×10^−4^, C_R_ = 0.3, μN = 0.25, N = 1.44×10^5^.

**Figure 2b:** 1000 replicate simulations with the following parameters: C_D_^0^ = 0.06, C_D_ = 2×10^−4^, C_P_ = 8×10^−5^, C_R_ = 0.35, μN = 0.025, N = 1.44×10^5^.

**Figure 2d: simulations with 1 antibiotic:** C_D_^0^ = 0.06, C_D_ = 2×10^−4^, C_P_ = 8×10^−5^, C_R_ = 0.35, N = 1.44×10^5^. μN = {0.1, 0.0875, 0.075, 0.0625, 0.05, 0.0375, 0.025, 0.0125, 0.00875}. 1000 replicate simulations per μN value. Each simulation was run until it reached an ESC-like state.

**simulations with 2 antibiotics:** C_D_^0^ = 0.06, C_D_ = 8×10^−4^, C_P_ = 4×10^−5^, C_R_ = 0.3, N = 1.44×10^5^. 1000 replicates per μN were run for 70000 ecological cycles for μN = {0.8, 0.7, 0.6, 0.5, 0.4, 0.3, 0.2}, 5000 replicates were run for 40000 ecological cycles for μN = 0.15. 5000 replicates were run for 160000 ecological cycles for μN = 0.1.

**Figure 3a:** C_D_^0^ = 0.06, C_D_ = 2×10^−4^, C_P_ = 8×10^−5^, C_R_ = 0.35, N = 1.44×10^5^, μN = 0.05.

**Figure 3b:** C_D_^0^ = 0.06, C_D_ = 8×10^−4^, C_P_ = 4×10^−5^, Cp0 = 0.3, N = 1.44×10^5^, μN = 0.4.

**Figure 3c:** C_D_^0^ = 0.06, C_D_ = 8×10^−4^, C_P_ = 4×10^−5^, C_R_ = 0.3, N = 1.44×10^5^, μN = 0.7.

**Figure 4a:** Number of antibiotics = 1. C_D_^0^ = 0.06, C_D_ = 2×10^−4^, C_P_ = 8×10^−5^, C_R_ = 0.35, N = 1.44×10^5^, μN = 0.05, m = {0.05, 0.01, 5×10^−4^, 5e-5, 1×10^−5^, 5×10^−6^, 1×10^−6^, 5×10^−7^, 1×10^−7^}, H = 10. 500 replicate simulations per μN. Each simulation was run until it reached an ESC-like state.

**Figure 4c:** C_D_^0^ = 0.03, C_D_ = 4×10^−5^, C_P_ = 4×10^−4^, C_R_ = 0.30, N = 1.44×10^5^, μN = 2, m = 5×10^−5^, H = 20.

**Supplementary Figure 1:** same as Figure 2d.

**Supplementary Figure 2:** same as Figure 2d.

**Supplementary Figure 3a: simulations with 1 antibiotic:** C_D_^0^ = 0.06, C_D_ = 2×10^−4^, C_P_ = 8×10^−5^, C_R_ = 0.35. N = {4.32×10^5^, 3.5×10^5^, 2.88×10^5^, 2.16×10^5^, 1.44×10^5^, 7.2×10^4^, 3.6×10^4^}, corresponding to μN = {0.1, 0.083, 0.0667, 0.05, 0.033, 0.01667, 0.00833}. 500 replicate simulations per μN were run until they reached an ESC-like state.

**simulations with 2 antibiotics**: C_D_^0^ = 0.06, C_D_ = 8×10^−4^, C_P_ = 4×10^−5^, C_R_ = 0.3. N = {3.6×10^5^, 2.88×10^5^, 2.16×10^5^, 1.44×10^5^, 7.2×10^4^}, corresponding to μN = {1, 0.8, 0.6, 0.4, 0.2}. 1000 replicate simulations per μN were run for 70,000 ecological cycles.

**Supplementary Figure 3b: simulations with 1 antibiotic:** C_D_^0^ = 0.06, C_D_ = 2×10^−4^, C_P_ = 8×10^−5^, C_R_ = 0.35, N = 1.44×10^5^, μN constant at 0.1 and N = {4.32×10^5^, 3.6×10^5^, 2.88×10^5^, 2.16×10^5^, 1.44×10^5^, 7.2×10^4^, 3.6×10^4^}. 1000 replicate simulations per μN were run until they reached an ESC-like state.

**simulations with 2 antibiotics:** C_D_^0^ = 0.06, C_D_ = 8×10^−4^, C_P_ = 4×10 ^−5^, C_R_ = 0.3, μN = 0.5 and N = {3.6×10^5^, 2.88×10^5^, 2.16×10^5^, 1.44×10^5^, 7.2×10^4^}. 1000 replicate simulations per μN were run for 70,000 cycles.

**Supplementary Figure 4:** C_D_^0^ = 0.06, C_D_ = 8×10^−4^, C_P_ = 4×10 ^−5^, C_R_ = 0.3, N = 1.44×10^−5^, μN = 0.5.

**Supplementary Figure 5:** C_D_^0^ = 0.06, C_D_ = 8×10^−4^, C_P_ = 4×10^−5^, C_R_ = 0.3, N = 1.44 × 10^−5^. {137, 121, 154} trajectories leading to community formation were classified by visual inspection for μN = {0.8, 0.4, 0.2}.

## Code Availability

The computer code is available from the corresponding author upon request.

## Data Availability

The datasets generated during this study are available from the corresponding author upon reasonable request.

## Supplementary Figures

**Supplementary Figure 1.**
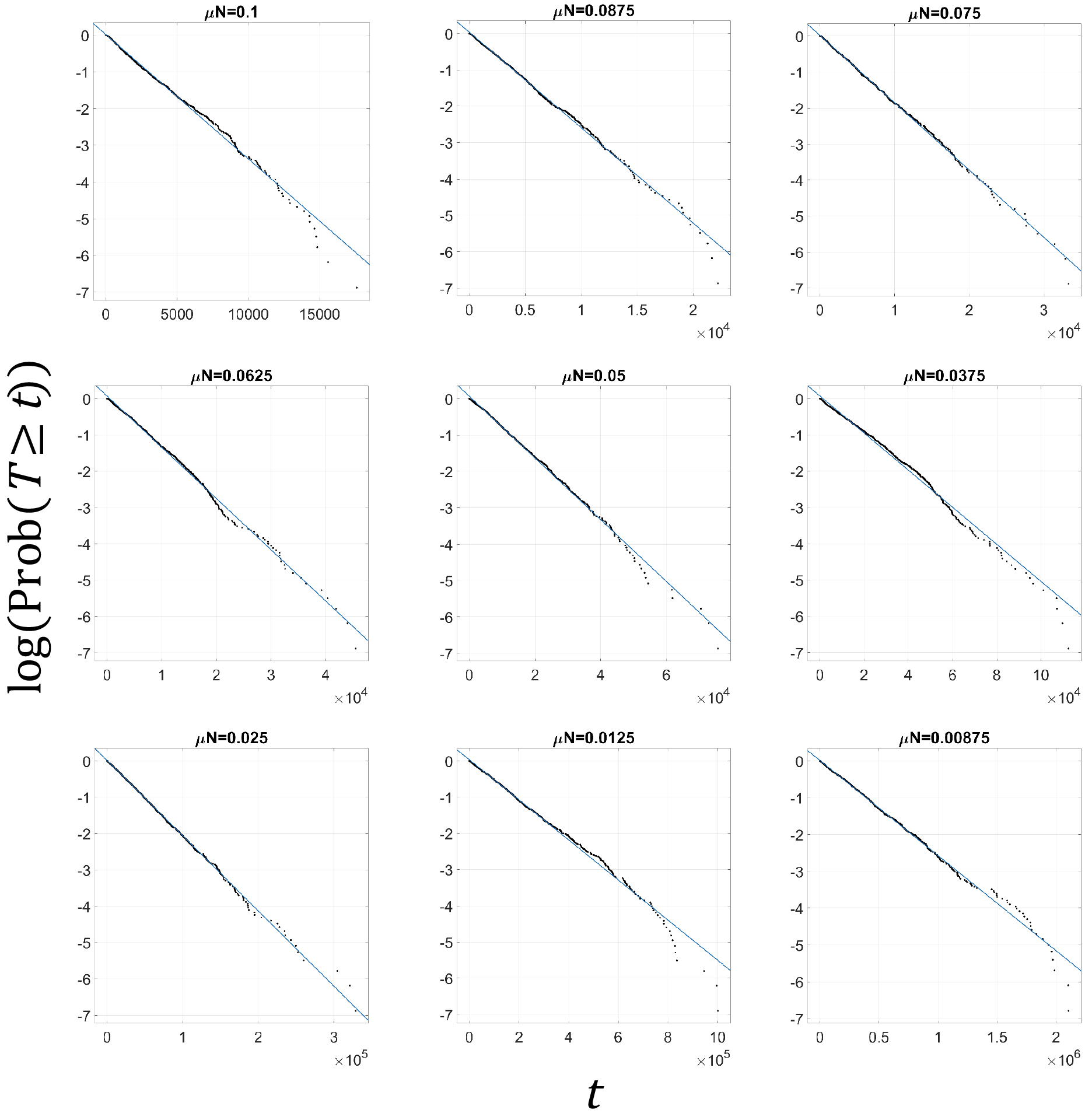
The community formation time, *T*, is exponentially distributed in simulations with 1 antibiotic. If *T* is exponentially distributed then Prob(*T* ≥ t) = exp(−rt). Shown are the empirically determined distributions of *T* for different *μN* (black points), demonstrating proportionality between log Prob(*T* ≥ *t*) and *t*. The slopes of the fitted lines (blue) provide an estimate of the rate of community emergence *r*.

**Supplementary Figure 2.**
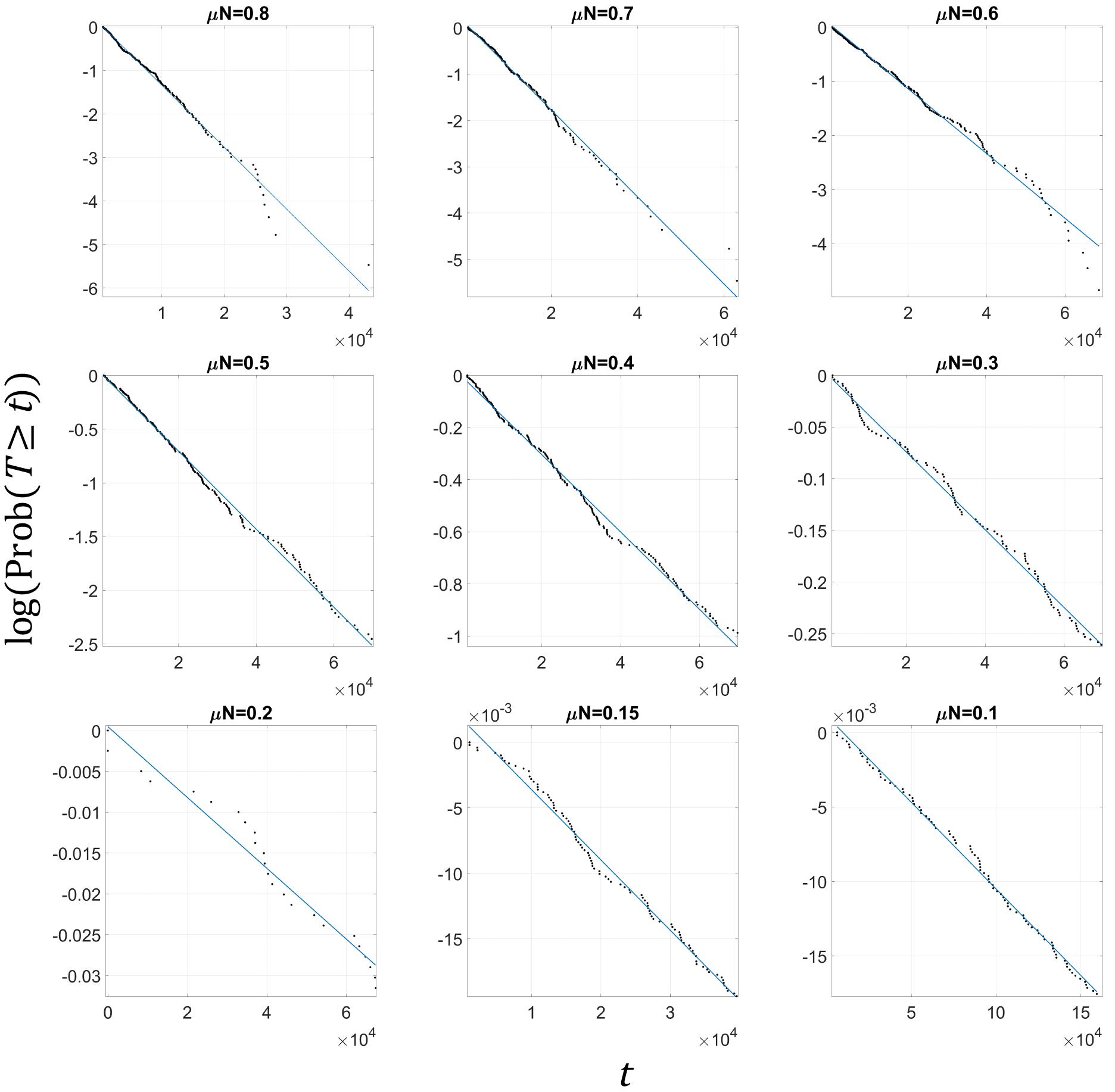
The community formation time, *T*, is exponentially distributed in simulations with 2 antibiotics. If *T*is exponentially distributed then Prob(*T* ≥ *t*) = exp(−*rt*). Shown are the empirically determined distributions of *T* for different *μN* (black points), demonstrating proportionality between log Prob(*T* ≥ *t*) and *t*. The slopes of the fitted lines (blue) provide an estimate of the rate of community emergence *r*.

**Supplementary Figure 3.**
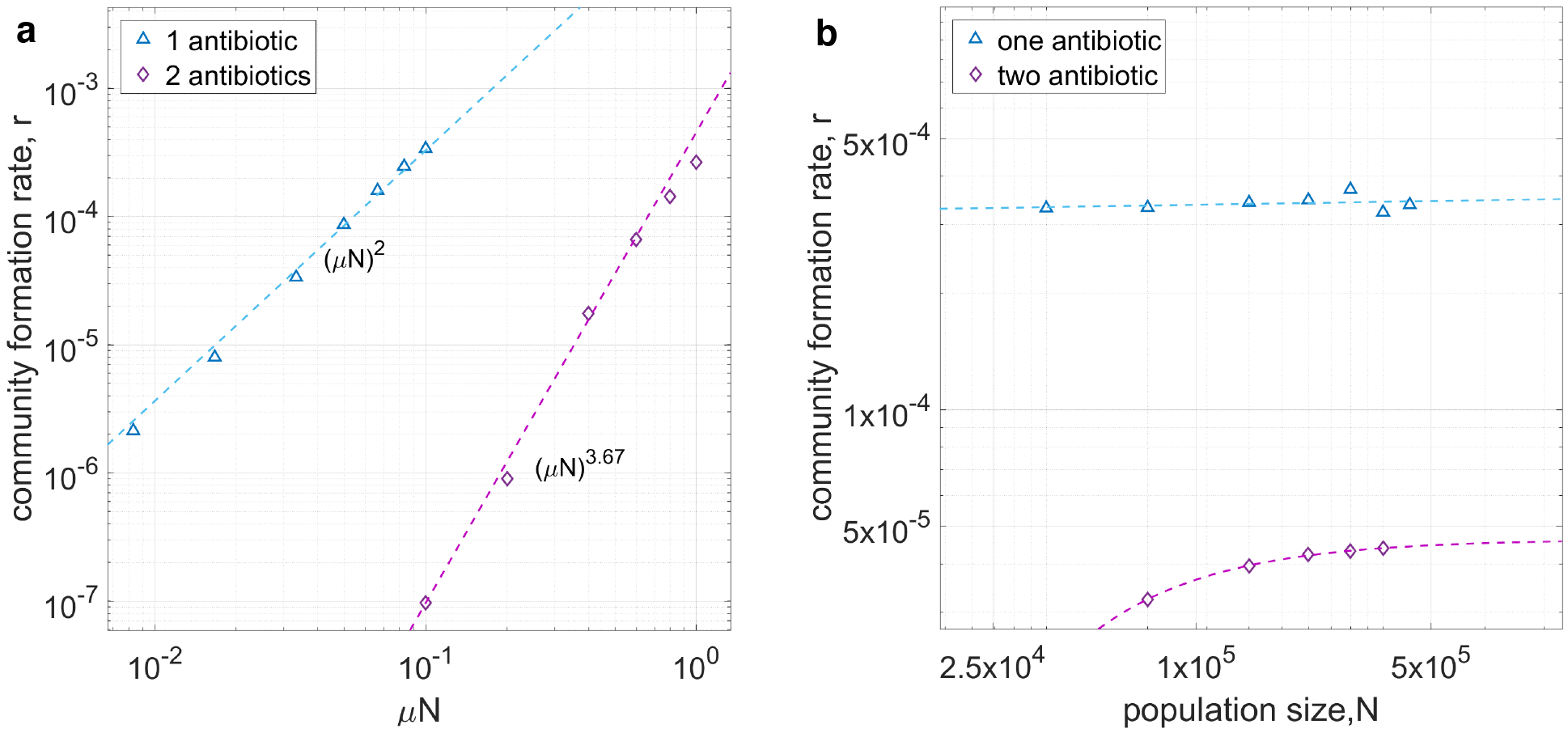
The community formation rate, *r*, is calculated for different values of *N* at fixed *μ* or *μN*. This complements Fig. 2d, where *μ* is changed at constant *N*. **(a)** *N* is changed at constant *μ* in simulations with one antibiotic (blue triangles) and two antibiotics (purple diamonds). The dotted lines are based on the scaling relationships found in Fig. 2d. The data reveals that *r* exhibits the same scaling with *μ* and *N*, indicating that *r*(*μ, N*) = *r*(*μN*). **(b)** *N* is changed at constant *μN*. For one antibiotic, *r* does not depend on *N* for the examined range. For two antibiotics, *r* depends weakly on *N* at low *N*, but becomes independent of *N* for sufficiently large populations. This data confirms that *r*(*μ, N*) = *r*(*μN*) for large populations. Therefore, the relevant control parameter is the number of mutations per population per ecological cycle *μN*.

**Supplementary Figure 4.**
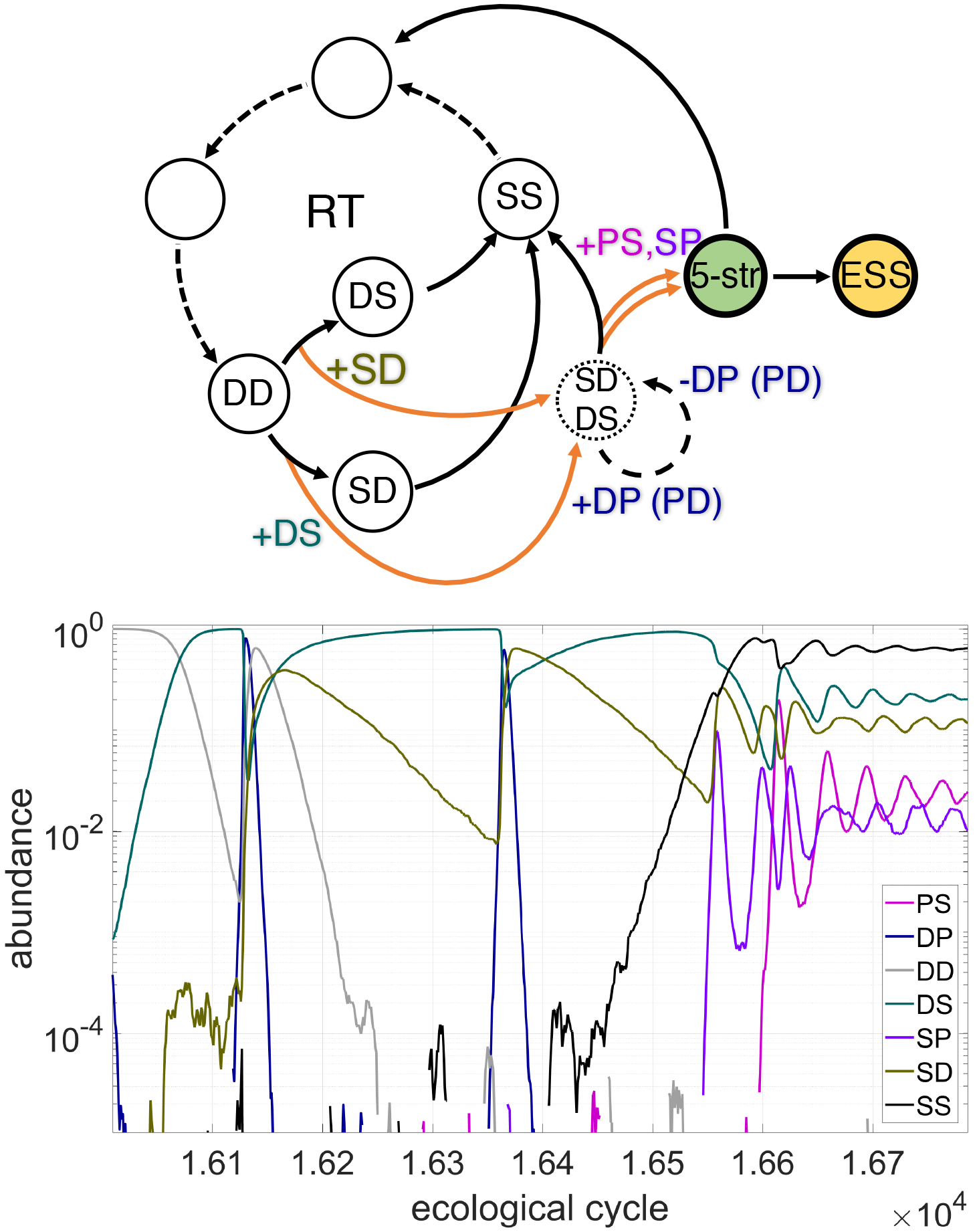
An alternative trajectory for 5-strain community formation in simulations with 2 antibiotics. This is a variant of the trajectory presented on Fig. 3c in which DS and SD strains persist for an extended time even though they are not close in fitness. DS can rapidly outcompete SD if the two strains are alone, but their coexistence is maintained by periodic invasions and extinctions of DP strains, which can kill DS when SD is low in abundance and cannot provide cross-protection.

**Supplementary Figure 5.**
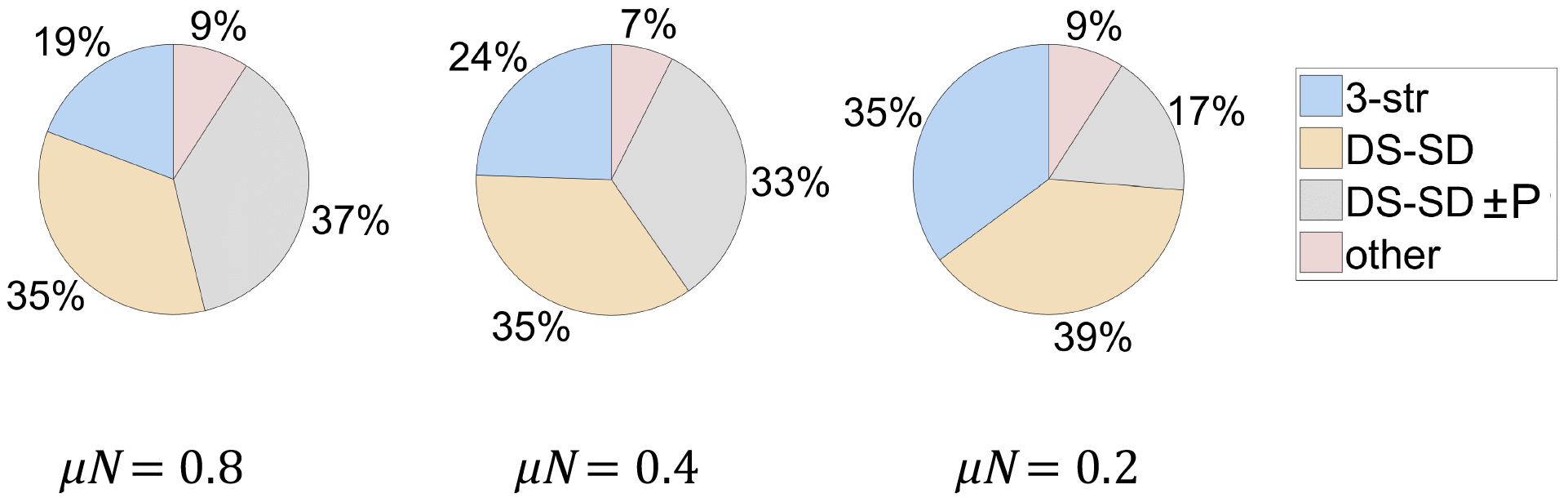
The majority of pathways for formation of a 5-strain community fall into 3 categories. Three main classes of pathways were observed: 1) trajectories that passthrough an ecologically stable intermediate community with 3 strains, as in Fig. 3b (blue, “3-str”); 2) trajectories that pass through a SD-DS 2-strain intermediate state that is unstable but persists because of near neutrality, as in Fig. 3c (yellow, “DS-SD”); 3) trajectories that pass through a SD-DS intermediate state where the two strains are substantially different in fitness, but persistence is prolonged by periodic invasions and extinctions of producer strains, as in Supplementary Fig. 4 (grey, “DS-SD±P”). Shown is the proportion of different classes for three different *μN*, as determined by visually scoring of trajectories leading to community formation. The proportion of trajectories that pass through a stepping-stone 3-strain community gradually increases with the decrease of *μN*. Strikingly, the power law scaling of *r* with *μN* is maintained despite the significant changes in the ratios of different channels of community formation.

